# Functional screening identifies novel miRNAs inhibiting Vascular Smooth Muscle Cell proliferation

**DOI:** 10.1101/2024.04.04.587890

**Authors:** Julie Rodor, Eftychia Klimi, Luca Braga, Nadja A.R. Ring, Margaret D. Ballantyne, Vladislav Miscianinov, Francesca Vacante, Katarina Miteva, Despoina Kesidou, Matthew Bennett, Abdelaziz Beqqali, Mauro Giacca, Serena Zacchigna, Andrew H. Baker

## Abstract

Proliferation of vascular smooth muscle cells (vSMCs) following injury is a crucial contributor to pathological vascular remodelling. MicroRNAs (miRNAs) are powerful gene regulators and attractive therapeutic agents. Here, we aim to systematically identify and characterise miRNAs with therapeutic potential in targeting aberrant vSMC proliferation. We performed a high-throughput *in vitro* screen using a library of 2042 human miRNA-mimics for their impact on vSMC proliferation and identified seven novel antiproliferative miRNAs i.e miR-323a-3p, miR449b-5p, miR-491-3p, miR-892b, miR-1827, miR-4774-3p, miR-5681b. Overexpression of these seven miRNAs affects proliferation of vSMCs from different vascular beds. Focusing on vein graft failure, a condition in which miRNA-based therapeutics can be applied to the graft ex-vivo, we showed that these miRNAs reduced human saphenous vein SMC (HSVSMC) proliferation without inducing apoptosis or senescence, and five of them also significantly decreased migration. HSVSMC transcriptomic analysis showed that each miRNA overexpression affects a core cell cycle gene network. However, this effect is mediated by distinct miRNA targets. In contrast to HSVSMC, miRNA overexpression in saphenous vein endothelial cells (ECs) led to no decrease or a less pronounced reduction in proliferation for the seven miRNAs. Transcriptomics analysis confirmed a distinct and limited response of ECs to the miRNA overexpression.

## INTRODUCTION

Vascular remodelling is an essential process of adaptive structural change in the vessel wall. It involves changes in vascular wall thickness, which confers elevated vascular resistance in response to pathological, haemodynamic, or iatrogenic injurious cues. However, this process can become aberrant, resulting in the development of vascular pathologies such as atherosclerosis, pulmonary hypertension^1^ as well as the intimal hyperplasia underlying saphenous vein graft failure. ^2^

Integral to the aetiology of the vascular remodelling process is the switch of resident vascular smooth muscle cells (vSMCs) from a differentiated and quiescent to a de-differentiated, pro-proliferative and pro-migratory phenotype.^3,4^ On the molecular level, excessive proliferation of vSMC is a complex process linked to the injury of the endothelial cell layer and the subsequent inflammatory response.^5^ The growth factor PDGF has been shown to play a crucial role in vSMC response^5^ and activation of PDGF in combination of IL1a pathway has been shown to promote proliferation *in vitro* in human saphenous vein SMC.^6^

Targeting vSMC proliferation is thus an attractive therapeutic strategy for preventing adverse vascular remodelling in response to injury. Approaches based on reducing vSMC proliferation have been successful pre-clinically and shown promising clinical results, as demonstrated by animal studies and clinical trials testing the use of anti-proliferative pharmacological agents in drug-eluting stents used for coronary angioplasty.^7^ The pitfall of these approaches, however, has been interference with re-endothelisation, a process integral in countering the subsequent pathological vascular remodelling events that initiate following endothelial cell injury and denudation.^7,8^ In the case of vein graft failure, therapy based on gene transfer has been considered and developed due to *ex vivo* access of the graft at the time of surgery.^9,10^

MicroRNAs (miRNAs) are small non-coding RNA molecules (20-24 nucleotides in length) that regulate gene expression through imperfect base-pairing with regions in the 3’UTR of target messenger RNA (mRNA), inducing their degradation or translational repression.^11^ MiRNAs have been shown to play critical roles in a range of biological contexts, including development, cancer, neurodegenerative disease.^12^ In cardiovascular physiology, dysregulation of several miRNAs has been implicated in the development of multiple diseases.^13^ For example, the SMC-enriched miR-143/145 cluster, has been involved in vascular remodelling across several diseases such as neointimal lesion formation, pulmonary arterial hypertension, and atherosclerosis.^14^ Multiple miRNAs regulating vSMC phenotypes^15^ and more particularly vSMC proliferation^16^ have also been reported.

Due to their potent effect in regulating multiple gene expression changes, miRNA based therapeutics approach has been developed in multiple disease contexts, including cardiovascular disease, with multiple ongoing clinical trials.^17,18^ These approaches rely on the ability to modulate miRNA abundance using miRNA mimics, inhibitors, or viral vector-mediated overexpression of miRNA loci. To select the best miRNA candidates for therapy in an unbiased way, functional screen has been developed. In particular, microscopy-based screen has been used successfully, initially using hundreds of miRNAs and more recently using a library of 2042 miRNAs, covering all miRNAs present in miRBase database.^19^ In the cardiovascular field, miRNA regulating cardiomyocyte proliferation have been identified^20^ and further testing showed that one of them, miR-199a, stimulates cardiac regeneration *in vivo*, in mouse^20^ and also in pig.^21^

Here, we implemented a miRNA high-content imaging screen to identify miRNA regulating vascular smooth muscle cell proliferation. Seven novel miRNAs with an anti-proliferative effect across SMC from different vascular bed were identified and further studied in the context of vein graft failure. In saphenous vein SMCs, we confirmed a strong decrease of proliferation, with no detrimental effect, upon overexpression of the 7 miRNAs. Transcriptomics analysis combined to miRNA target prediction showed distinct targets for the 7 miRNAs, leading to a common regulation of a network of core cell cycle genes. In contrast, we observed limited proliferation and transcriptomics changes in saphenous vein endothelial cells after miRNA overexpression.

## RESULTS

### Functional screening identifies miRNAs that block vSMC proliferation

To identify miRNAs exerting an antiproliferative phenotype in vSMCs in an unbiased manner, we performed a high-throughput high content miRNA screening in human pulmonary artery SMCs (HPASMC) using a library of miRNA mimics corresponding to 2042 unique human mature miRNA sequences (Figure 1A). The effect on proliferation was assessed based on EdU incorporation. High-content fluorescent image analysis was used to quantify viable (Hoechst33342+) and proliferating (EdU+/Hoechst33342+) HPASMCs. The screen included the overexpression of four different miRNA controls (miR-CTRL) and effect on proliferation was assessed relative to the average of the 4 miR-CTRL (Supplementary Table 1).

**Figure 1.**
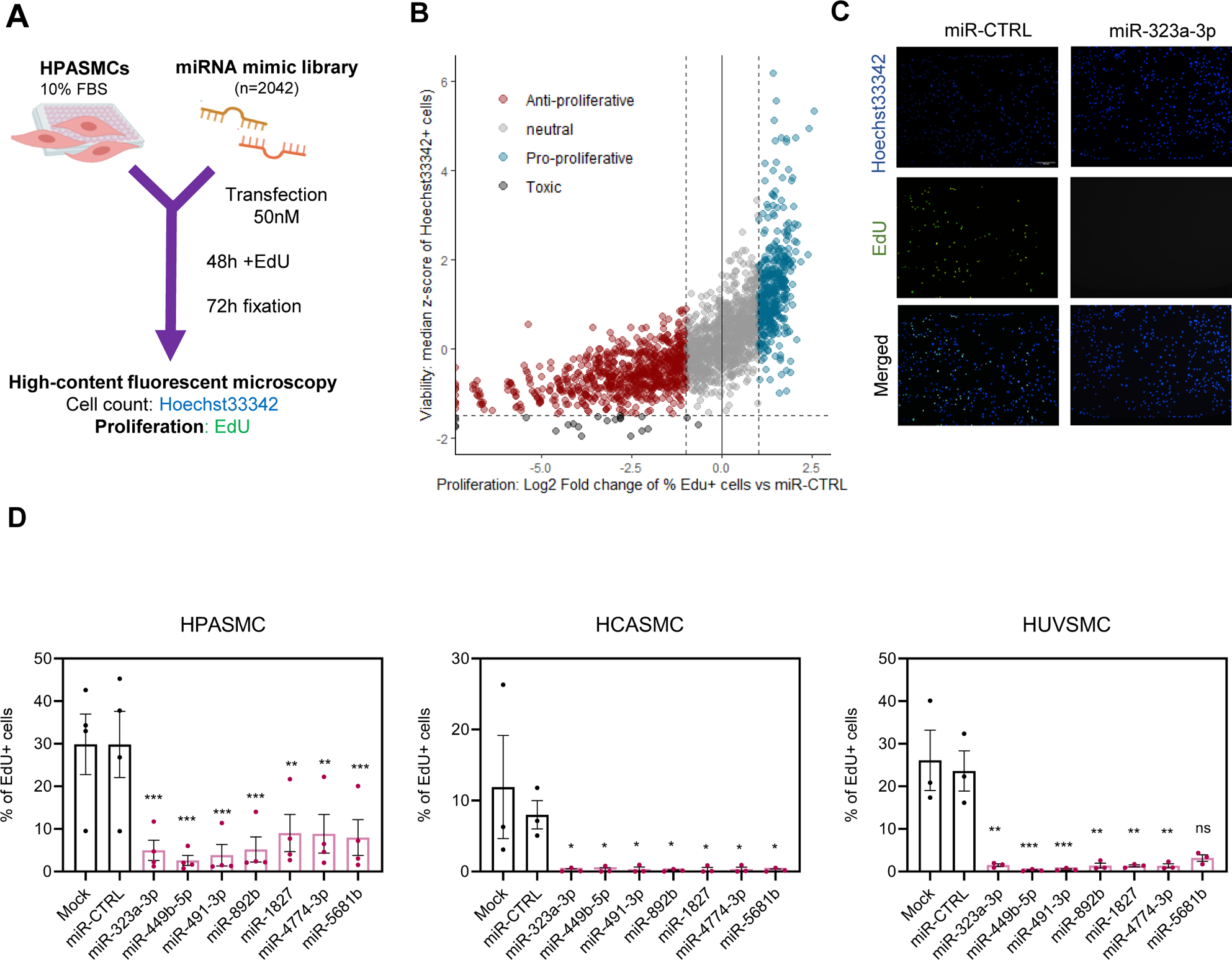
High-throughput miRNA screen identifies novel miRNAs regulating VSMC proliferation. A. Schematic of high-throughput miRNA screen design in vSMC. B. Scatter plot of viability (expressed as median z-score of Hoechst3342+ cells) and proliferation (expressed as Log2 fold change of %EdU+ cells vs miR-CTRL) changes for each mimic-mediated miRNA overexpression (n=1). C. Representative images of human PASMCs stained with Hoechst3342 (blue) and EdU (green) following treatment with miR-323a-3p mimic and miR-CTRL. Scale bar is 100μm. D. High-throughput microscopy imaging quantification of EdU incorporation in (D) HPASMCs (n=4), (E) HCASMCs (n=3) and (F) HUVSMCs (n=3) transfected with 7 candidate miRNA mimics or miR-CTRL, as well as the “mock” transfection control. Statistical analyses were performed using Iman and Conover non-parametric ranking followed by a repeated-measure ANOVA and the P-value was calculated for the comparison between miRNA mimic treatment and miR-CTRL using Dunnett’s test for multiple comparisons. On the graphs, \**P* < 0.05, \*\**P* < 0.01, \*\*\**P* < 0.001, ns: non-significant. n numbers correspond to distinct biological replicates.

First, 22 miRNAs with a large decrease on cell count (Hoechst33342+) were discarded from our analysis due to likely toxicity (Figure 1B). Based on a 2-fold change in the percentage of EdU+ cells relative to the 4 miR-CTRL, we identified 715 miRNAs with an anti-proliferative effect and 417 miRNAs with a pro-proliferative effect in PASMCs (Figure 1B and Supplementary Table 1). The list of miRNAs included several miRNAs with known effects on vSMC proliferation^16^, providing confidence on the screen result validity. Indeed, the known pro-proliferative miRNA miR-146a and miR29a had a positive effect on HPASMC proliferation in the screen assay while the known anti-proliferative miRNAs miR-124, miR-214 and miR-34a showed a decrease of HPASMC proliferation (Supplementary Table 1).

Ten miRNAs had a strong effect on PASMC proliferation, reducing the number of EdU+ nuclei to 0 without a marked reduction in cell count (Supplementary Table 1). The list of 10 miRNAs included miR-34a-3p, a known anti-proliferative miRNA mentioned above. We selected 7 miRNAs for further study based on novelty in the cardiovascular field: miR-323a-3p (Figure 1C), miR-449b-5p, miR-491-3p, miR-892b, miR-1827, miR-4774-3p and miR-5681b (Supplementary Figure 1).

The effect of mimic-mediated overexpression of these 7 miRNAs on PASMC proliferation was confirmed on 4 additional biological replicates (Figure 1D). We also tested the effect of miRNA overexpression on smooth muscle cells from different vascular beds using the same conditions and technology as the screen. All 7 novel miRNA candidates showed a significant reduction in EdU incorporation in human coronary artery smooth muscle cells (HCASMCs) (Figure 1E). In human umbilical vein smooth muscle cells (HUVSMCs), 6 out of the 7 miRs (miR-323a-3p, miR-449b-5p, miR-491-3p, miR-892b, miR-1827, miR-4774-3p) showed a significant decrease in the percentage of EdU+ cells (Figure 1E).

This data suggests that the overexpression of these 7 novel miRNAs could have therapeutic potential to reduce vSMC proliferation across different vascular pathologies.

### Overexpression of miRNA candidates reduces proliferation and migration of HSVSMCs, without inducing apoptosis or senescence

As SMC proliferation contributes to pathological remodeling in vein graft failure and therapeutic intervention can be implemented ex-vivo at the time of grafting, miRNA-based therapy could be a suitable strategy for this pathology. Hence, we investigated further the expression and effect of the 7 miRNAs in primary human saphenous vein SMCs. We performed mimic-mediated overexpression of the 7 miRNAs and HSVSMC phenotypic switching (de-differentiated, pro-proliferative and pro-migratory phenotype) was induced using a combination of IL1-α and PDGF-BB as previously described ^6^ (Figure 2A). We assessed the effect of the miR overexpression on proliferation and migration but also checked for any potential detrimental effect by analysing apoptosis and senescence (Figure 2A).

**Figure 2:**
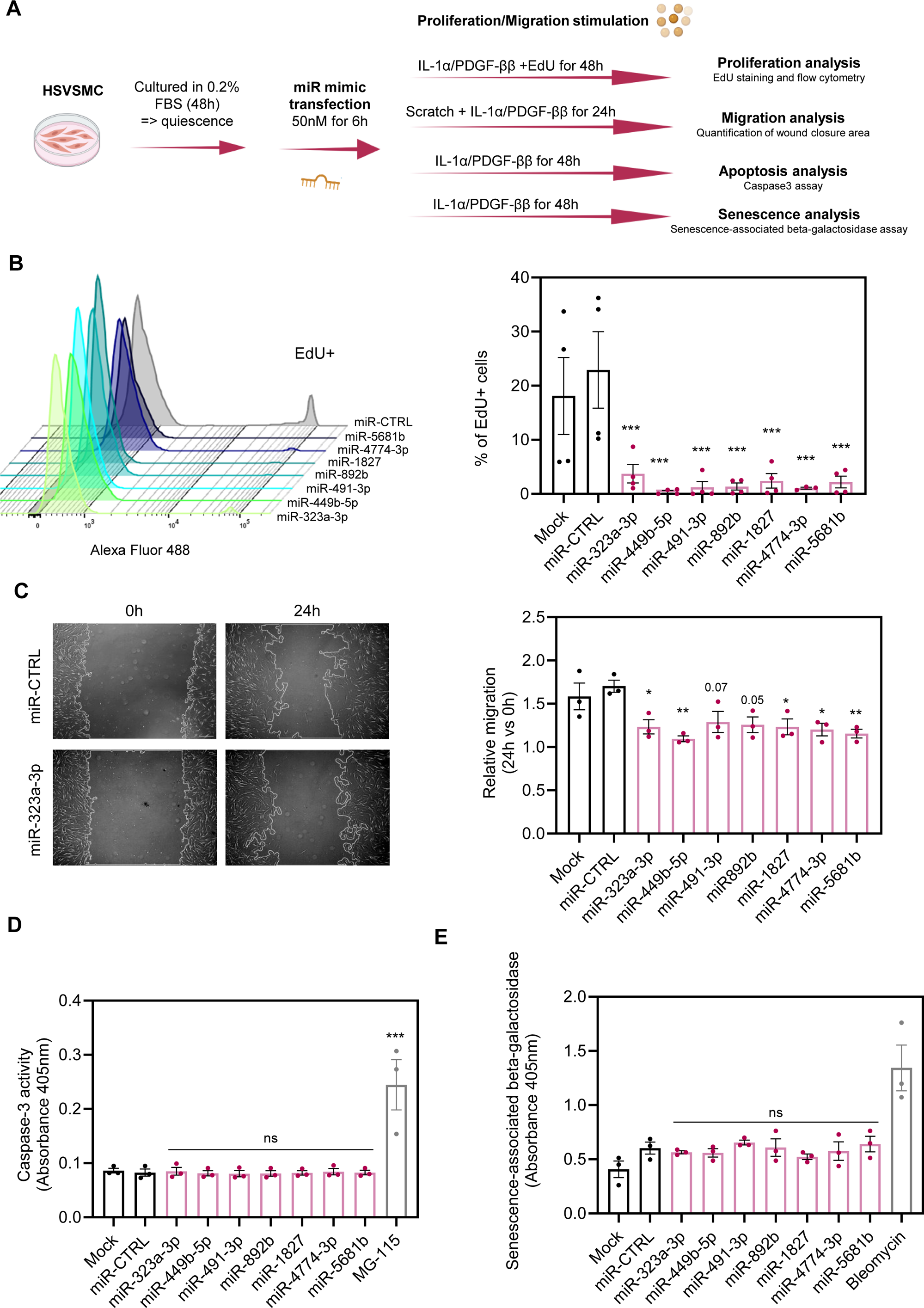
The overexpression of 7 novel candidate miRNA affects HSVSMC proliferation and migration *in vitro*. A. Schematic of experimental design for assessing the effect of miRNA overexpression on IL-1α/PDGF-BB treated HSVSMCs. B. Flow cytometric quantification of EdU incorporation in IL-1α/PDGF-BB -stimulated HSVSMCs transfected with the 7 miRNA mimics vs miR-CTRL and Lipofectamine treated cells (Mock). On the left, representative EdU plot and on the right, bar graph quantification (n=4, except n=3 for miR-4774-3p). Statistical analyses were done using a Mixed-Effects model. C. Wound healing assay of IL-1α/PDGF-BB-stimulated HSVSMCs transfected with the 7 miRNA mimics or miR-CTRL, as well as the “mock” transfection control. On the left, representative images from scratch assay at 0- and 24-hours for miR-CTRL and miR-323a-3p. Scale bar is 500µm. On the right, quantification of wound healing (24h area versus 0h area) via the ImageJ MRI wound healing tool (n=3). Statistical analyses were done using Iman-Conover non-parametric ranking followed by RM ANOVA D. Quantification of Caspase-3 activity (absorbance measured at 405nm) in IL-1α/PDGF-BB - stimulated HSVSMCs transfected with the 7 miRNA mimics or miR-CTRL, as well as the “mock” transfection control (n=3). The proteasome inhibitor MG-115 was used as a positive control for apoptosis induction. Statistical analyses were done using Iman-Conover non-parametric ranking followed by RM ANOVA. E. Quantification of Senescence Associated (SA) β-galactosidase (β-gal) activity (absorbance measured at 405nm) in IL-1α/PDGF-BB -stimulated HSVSMCs transfected with the 7 miRNA mimics or miR-CTRL, as well as the “mock” transfection control (n=3). Bleomycin (1µg/mL) was used as a positive control for senescence induction. Statistical analyses were done using Iman-Conover non-parametric ranking followed by RM ANOVA. P-values for the comparison between miRNA mimic treatment and miR-CTRL treatment obtained after Dunnet’s test for multiple corrections are included on the graph: \**P* < 0.05, \*\**P* < 0.01, \*\*\**P* < 0.001, ns= non-significant. n numbers correspond to distinct biological replicates.

Overexpression of the 7 miRNAs was confirmed by RT-qPCR (Supplementary Figure 2). It was noted that all 7 miRNAs have low or no endogenous expression (based on qPCR cycle threshold value - data not shown) in quiescent and IL1a/PDGF-BB-treated HSVSMCs suggesting that any phenotype observed in mimic-treated HSVSMCs will be linked to their exogenous expression. Flow cytometric quantification of EdU incorporation in HSVSMCs showed significant decreases in IL-1α/PDGF-BB-induced proliferation (decrease ranging from 83.7 to 98.2%) after transfection with the seven miRNA mimics compared to miR-CTRL (Figure 2B). We also observed that 5 out of the 7 miRNAs significantly reduced the migration rate of IL-1α/PDGF-BB-stimulated HSVSMC, assessed by scratch wound assay (Figure 2C and Supplementary Figure 3). No change in caspase-3 activity was detected upon the overexpression of the 7 miRNAs in HSVSMCs, in contrast to the increase of caspase-3 activity observed with the treatment with the proteasome inhibitor MG-115 used as a positive control (Figure 3C). This data shows that none of the miRNAs induce apoptosis in HSVSMCs. Furthermore, no increase in senescence, measured by senescence-associated (SA) β-galactosidase activity, was observed following overexpression of the 7 miRNAs (Figure 3D).

**Figure 3:**
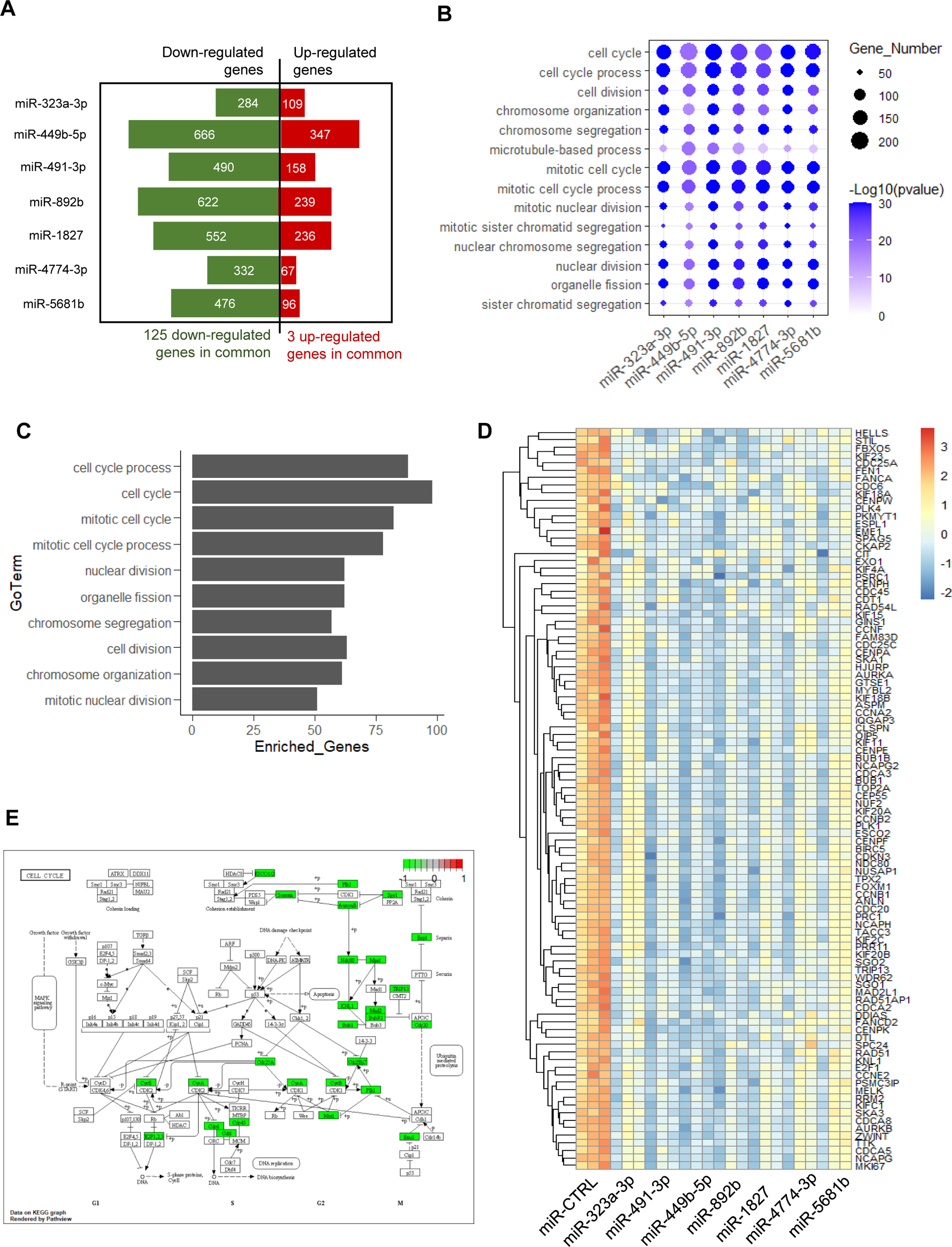
Transcriptomic changes upon overexpression of the 7 candidate miRNAs in HSVSMCs. A. Number of significant differentially expressed genes for each miRNA overexpression versus miR-CTRL. The number of genes commonly regulated by all miRNAs is indicated below. B. Plot of the Top10 enriched GoTerms identified for the genes regulated by each miRNA overexpression. C. Enriched GoTerms for the 125 genes commonly down-regulated by all 7 miRNAs. D. Heatmap of the 98 cell-cycle genes commonly down-regulated by all 7 miRNAs. E. Diagram showing the genes regulated by all 7 miRNAs (in green) involved in cell cycle KEGG pathway (hsa04110).

These results demonstrate that overexpression of the candidate miRNAs reduces IL-1α/PDGF-BB-induced HSVSMCs proliferation and migration, without causing HSVSMC apoptosis nor senescence, making them good candidate for therapy in vein graft failure context.

### Transcriptome analysis revealed the regulation of a common network of cell cycle genes by all 7 miRNAs

To understand the effect of the miRNA overexpression at the transcriptomic level, we performed RNA sequencing (RNAseq) on quiescent HSVSMC and IL-1α/PDGF-ββ induced HSVSMC treated with the different miRNA mimics. The principal component analysis (PCA) showed distinct clusters based on conditions and patient origins of the HSVSMCs (Supplementary Figure 4A). As expected, IL-1α/PDGF-ββ induced HSVSMC clustered separately from quiescent cells (Supplementary Figure 4A). After removal of batch/patient effect, the PCA analysis clearly showed that all miRNAs treated samples did not overlap with their relative IL-1α/PDGF-ββ controls (no treatment, mock treatment or miR-CTRL) and were also distinct from quiescent cells (Supplementary Figure 4A). This revealed that miRNA overexpression affects the transcriptome but does not revert it to a “quiescent cell” transcriptome. We performed a differential expression analysis between each miRNA mimic overexpression condition and miR-CTRL condition. As we observed sample separation based on patient origin in the PCA plot (Supplementary Figure 4A), we corrected for patient variance in the differential gene expression analysis. Based on a two-fold change threshold, we detected between 393 and 1013 differentially expressed genes due the miRNA overexpression (Figure 3A and Supplementary Table 2). For each list of differentially expressed genes, we performed Gene Ontology (GO) analysis (Supplementary Table 3) and found that the top 10 enriched GO Terms for each miRNA overexpression heavily overlapped between the 7 miRNAs and were exclusively related to cell cycle process (Figure 3B). This is in agreement with the strong effect of the miRNA overexpression on HSVSMC proliferation and shows that similar pathways are regulated by the 7 miRNAs. As some of the miRNA overexpression also led to a migratory phenotype, we screened the list of enriched Go-terms and found migration-related GO Terms for miR-449b-5p, miR-491-3p, miR-892b, miR-1827, miR-4774-3p and miR-5681b (Supplementary Table 3). As the regulation of similar pathways by the 7 miRNAs might correspond to the regulation of similar genes, we assessed the overlap of the differentially regulated genes and identified 3 genes commonly up-regulated and 125 genes commonly down-regulated by all 7 candidate miRNAs (Figure 3A). Interestingly, the enriched GO Terms of the 125 down-regulated genes were related to cell cycle process (Figure 3C). We found 98 out of the 125 down-regulated associated with the “Cell Cycle” GO Term annotation (Figure 3D) and 17 of them were associated with the KEGG cell cycle pathway which includes the main components required for mitotic cell cycle progression (Figure 3E). This analysis suggests the anti-proliferative phenotype is mediated by the regulation of the same core cell cycle genes and cell cycle regulators for all seven miRNAs.

### The mechanism of action of the novel miRNA involved distinct targets

In order to characterise the mechanism of action of each miRNA, we aimed to identify their direct targets in proliferating HSVSMCs. As miRNAs are negative post-transcriptional regulators of gene expression, we focused our analysis on down-regulated genes upon miRNA overexpression and used prediction tools to identify potential miRNA-mRNA interaction (Figure 4A). We used multimiR, a prediction tool package that compiles several target prediction algorithms ^22^, and kept predicted targets ranked in the top 50% of at least two prediction tools. We identified between 13 and 224 candidate targets for the different miRNAs (Figure 4B and Supplementary Table 4), with a total of 680 candidate targets if we consider all 7 miRNAs together. 89.4% of the identified 680 targets are unique to one specific miRNA while only 72 genes (10.6%) were candidate targets for at least 2 miRNAs (Figure 4C). We did not identify any gene targeted by all 7 miRNAs but revealed that IGF2BP3 could be targeted by 5 of the 7 miRNAs. This data suggests that each miRNA have distinct mechanism of action. Interestingly, miR-323a-3p, miR-449b-5p, miR-892b and miR-1827 can directly target some of the 125 common down-regulated genes, suggesting a possible direct mechanism on this network while the regulation of the network by the other miRNAs might be indirect. Then, we assessed if any of the 7 miRNA candidate target genes could be involved in the process, progression and/or regulation of cell cycle using the GO Term annotation “Cell Cycle”. We found that 6 out of the 7 miRNAs have at least one target gene related to “Cell Cycle”” though these are still mostly different candidate targets per miRNA (Figure 4D).

**Figure 4:**
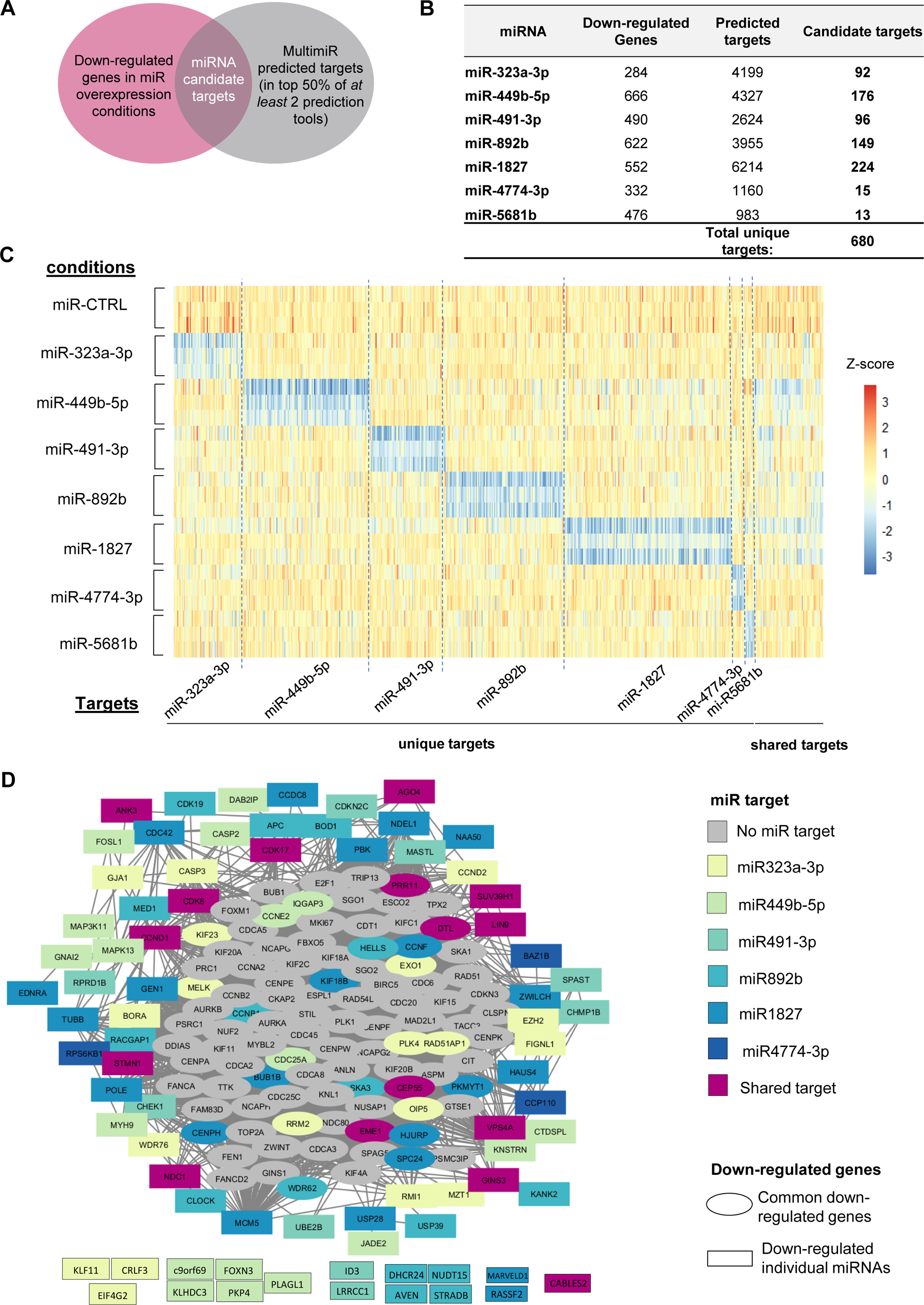
The 7 candidate miRNAs regulate distinct targets. A. Approach to identify miRNA targets. B. Number of down-regulated genes, predicted targets and candidate targets for each miRNA. C. Heatmap of all candidate targets for the 7 miRNAs in the HSVSMC RNAseq with a separation between unique and shared targets. D. Network of candidate targets and commonly down-regulated involved in Cell Cycle. The predicted gene interaction was obtained using STRING. The network was visualised using Cytoscape. The commonly down-regulated genes are represented in the centre of the network while the targets are located at the periphery.

We also performed a Gene Ontology analysis of the miRNA candidate targets (Supplementary Figure 5). MiR-4774-3p and miR-5681b were not included in this analysis as their number of candidate targets is too low (15 and 13 candidates respectively). We only found cell-cycle enriched GO Terms for the targets of miR-323a-3p (Supplementary Figure 5A) while other processes were enriched for the other miRNA targets. Interestingly, migration-related terms were observed in the analysis of miR-449b-5p targets (Supplementary Figure 5B), which coincided with miR-449b-5p overexpression having the strongest decrease of HSVSMC migration among the 7 novel miRNAs. For miR-892b, terms related to osteoblast differentiation, another SMC phenotype linked to vascular disease ^23^, were identified (Supplementary Figure 5D).

Our target analysis revealed each miRNA have different candidate targets. We identified some targets relevant to cell cycle regulation, suggesting they might have a driving role in the anti-proliferative phenotype, but we also highlighted the enrichment of other processes.

### Overexpression of individual miRNA candidates differentially regulates HSVSMCs and HSVECs

Because preservation of the function of the endothelium during vein grafting is key, we overexpressed the 7 miRNAs in human saphenous vein endothelial cells (HSVECs) and assessed the effect on proliferation based on EdU incorporation and flow cytometry (Figure 5A). No significant change on proliferation was observed for miR-323a-3p, miR-491-3p, miR-892b, miR-1827 and miR-5681b compared to miR-CTRL (Figure 5B). A significant decrease in proliferation was observed for miR-449b-5p and miR-4774-3p. However, while a 98% and 95% decrease of proliferation was observed in HSVSMC for miR-449b-5p and miR-4774-3p respectively, the effect on endothelial cell was lower with only a 61% decrease of proliferation for both miRNAs.

**Figure 5:**
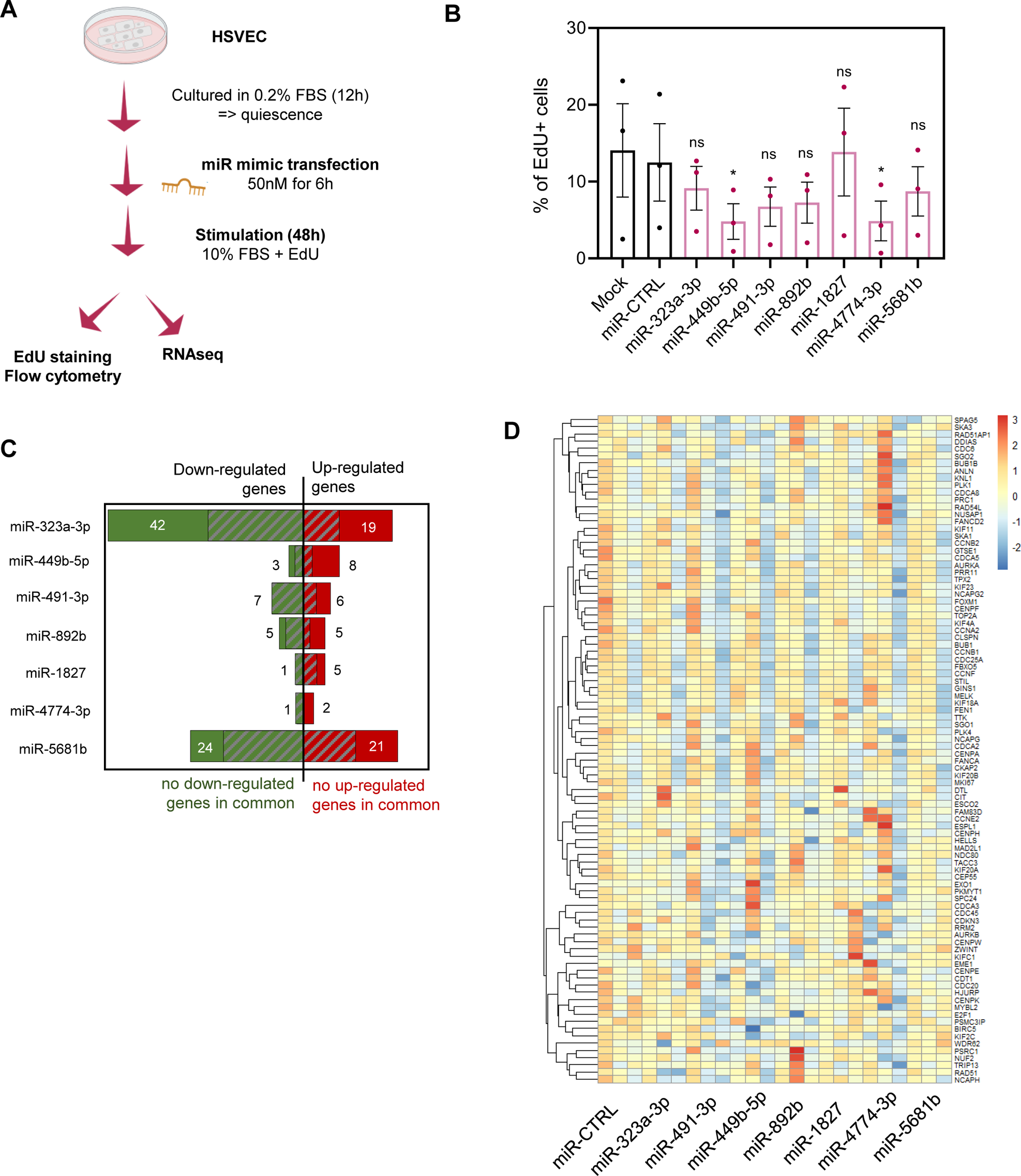
Proliferation and transcriptomic changes upon overexpression of the 7 candidate miRNAs in HSVECs. A. Schematic of experimental design for assessing the effect of miRNA overexpression on HSVEC proliferation and transcriptome. B. Flow cytometric quantification of EdU incorporation in HSVECs transfected with the 7 miRNA mimics vs miR-CTRL and Lipofectamine treated cells (Mock). (n=3). Statistical analyses were done using Iman-Conover non-parametric ranking followed by RM ANOVA. The P-value was calculated for the comparison between miRNA miRNA mimic treatment and miR-CTRL using Dunnett’s test for multiple comparisons. On the graph, \**P* < 0.05, ns: non-significant. C. Number of significant differentially expressed genes for each miRNA overexpression versus miR-CTRL based on RNAseq in HSVECs. The number of genes commonly regulated by all miRNAs is indicated below. Grey areas show the proportion of genes also regulated in HSVSMCs. D. HSVEC expression profile of the cell-cycle genes commonly down-regulated by all 7 miRNAs in HSVSMC. 92 out of the 98 genes were detected in HSVECs.

To understand the effect of the 7 miRNA overexpression at a molecular level, we performed RNAseq. Contrary to what was observed in HSVSMC (Supplementary Figure 4), the principal component analysis did not reveal a clear separation between the miRNA overexpression condition and the miR-CTRL in HSVEC (Supplementary Figure 6), suggesting no or low effect of the miRNA overexpression on the transcriptome. Differential gene expression analysis identified changes for 3 to 61 genes across the different conditions with no gene commonly regulated across the conditions (Figure 5C and Supplementary Table 5), confirming a low effect on the transcriptome. As expected, some of the changes in ECs were previously identified in the HSVSMC RNAseq (Figure 5C), showing some common but minimal gene regulations by the miRNAs across these two cell types.

We also visualised the expression of the 98 genes commonly down-regulated across the overexpression of the miRNAs in HSVSMC and involved in cell cycle within the HSVEC RNAseq. We observed little difference between miRNA overexpression conditions and miR-CTRL for these genes in HSVEC (Figure 5D), in agreement with the lack or low proliferation phenotype in this cell type. Equally, we visualised the expression of the miR candidate targets identified in the HSVSMC RNAseq. 32 of the 680 miRNA candidate targets had low or no expression in HSVEC and the 648 expressed targets do not display a consistent down-regulation across the three replicates in HSVEC samples (Supplementary Figure 7).

This data indicates that the miRNAs can strongly regulate HSVSMCs without significantly affecting the proliferation and transcriptome of endothelial cells.

## DISCUSSION

Therapy targeting vascular smooth muscle cell proliferation could prevent vascular remodelling across different diseases. Here, we considered the use of miRNA-based therapeutics and performed a high-throughput functional miRNA screen to unbiasedly identify miRNAs regulating vascular smooth muscle cell proliferation. We focused our study on seven novel anti-proliferative miRNAs and confirmed their effect in proliferation in different types of vSMCs. Their overexpression in HSVSMC lead to decrease proliferation, and also migration for 5 of them, without any effect on apoptosis and senescence, suggesting their potential in the context of vein graft failure. Transcriptomic analysis showed the regulation of a core network of cell cycle genes for all seven miRNAs in agreement with the anti-proliferative phenotype. However, target analysis suggested distinct targets for each individual miRNA. Overexpression of the 7 miRNAs shows no or little effect on proliferation and gene expression changes in HSVEC, suggesting a HSVSMC specific effect and an added potential for therapy.

Our study is based on a high-throughput screen using a library of 2042 miRNAs corresponding to all miRNAs annotated in miRBase at the time of the study design. Therefore, it constitutes a comprehensive resource of miRNA overexpression effects in vSMC proliferation and we identified 715 miRNAs with a negative effect on PASMC proliferation, providing a large number of candidates for further studies. The 7 selected miRNAs, based on stringent criteria (i.e overexpression leading to 0 Edu+ cells), showed an anti-proliferative phenotype on vSMC from different vascular bed, suggesting their modulation could be beneficial in different disease contexts. This has been shown to be the case for several miRNAs involved in vSMC phenotypic changes.^14^ For example, miRNA-21 ablation was shown to attenuates neointima formation in a mouse model of vein graft remodelling^24^ while miR-21 modulation had also an impact in the context of abdominal aortic aneurysm.^25^ MiR-34a was found among the screen top anti-proliferative miRNAs and was previously implicated in several cardiovascular diseases and considered as an promising therapeutic candidates.^26^ However, miR-34a also regulates vascular senescence and inflammation^27^ processes that might be detrimental in some specific disease context.

The transcriptomic analysis in HSVSMC showed that all 7 miRNA overexpression lead to the down-regulation of the same network of cell cycle genes, in agreement with the shared anti-proliferative phenotypes. To provide miRNA candidate targets with high confidence, we incorporated the list of down-regulated genes in addition to a combination of several prediction tools. With this approach, we showed distinct target pools for each miRNA with no common target. IGF2BP3 was identified as a candidate target for 5 miRNAs. Interestingly, IGF2BP3 promotes cell proliferation in cancer^28^ but its role has not been investigated in vascular SMCs. We also found that six out of the seven miRNAs have candidate targets involved in cell cycle based on GO Term annotation and their regulation could directly explain the anti-proliferative phenotype. Interestingly, EZH2, a cell-cycle annotated gene and candidate target of miR-323-3p, was involved in the pro-proliferative phenotype of vSMC in PAH.^29^ In addition, knockdown of GNAI2, a target of miR-449b-5p, decrease the hyperproliferation of SMC in spontaneously hypertensive rats.^30^

While the regulation of individual cell cycle gene by each miRNA could be responsible for the anti-proliferative phenotype, each miRNA also targets different processes that could converge towards an anti-proliferative effect. Enrichment of “migration” related terms was found for miR-449b-5p targets and we know that pathological vSMC display both a pro-migratory and pro-proliferative phenotype with some connections between the two phenotypes.^31^ For miR-892b, we found an enrichment of “negative regulation of osteoblast differentiation” term for its targets. Recently, single cell RNAseq has revealed the diversity of differentiated vascular smooth phenotype in healthy and disease condition, with the identification of osteogenic-like SMC.^23^

As we do not want a miRNA-based strategy to have a detrimental effect on the endothelium, we assessed the effect of the miRNA overexpression on HSVEC and showed no significant or a limited effect on proliferation as well as gene expression changes. Therefore, a therapeutics approach using mimics, and thus not relying on a SMC-specific delivery, could work for these miRNA candidates. Cell-type specific effect of miRNAs have been previously reported^32^ but are poorly understood. The different response is believed to be linked to different target availability due to distinct transcriptome in each cell type. The miRNA candidate target identification was based on the HSVSMC RNAseq. 32 of these targets are not expressed in HSVEC and this difference could explain the difference in phenotype. For example, pregnancy-associated plasma protein A (PAPPA) gene is a candidate target for miR-892b but its expression is very low (less than 0.5 FPKM) in HSVEC and therefore not regulated in this cell type. Interestingly, PAPPA was shown to be regulated by miR-141^33^ and miR-490-3p^33^ in vSMC and contributing to vSMC proliferation.

Our *in vitro* study led to promising human miRNA candidates to take forward for therapeutics. Testing this candidate *in vivo* in different animal models would be the next important step but will depend on the conservation of the miRNA in the studied species or at least the conservation of the respective miRNA targets. Large animal models such as pig are particularly relevant for vein graft failure as the setting is similar to the human graft (saphenous vein interposition into carotid artery) and the vessel wall more comparable to human than any small animal models.^34^ Human ex-vivo vein graft model, modelling vascular remodeling^35^ could also been used to test the therapeutic potential of the miRNA overexpression. While mimic miRNA have been used in therapy settings,^18^ other expression and delivery methods, viral or non-viral^17^ might be necessary in vein graft settings to allow a targeted delivery to specific cell type or subtypes.

## MATERIALS & METHODS

### Cell Culture

Human pulmonary artery (HPASMCs), coronary artery (HCASMCs) and umbilical vein (HUVSMCs) smooth muscle cells were purchased from Lonza and cultured using the recommended media. Primary human saphenous vein derived smooth muscle cells (HSVSMCs) and endothelial cells (HSVECs) were isolated from medial explants. All donated tissues have been obtained under proper informed consent and the investigation conforms with principles in the Declaration of Helsinki. HSVSMCs were obtained and maintained as previously described ^6^. Briefly, HSVSMC cell culture was performed using Smooth Muscle Cell (SMC) growth medium 2 (Promocell) were supplemented with the SMC 2 medium supplement (Promocell), 10% foetal bovine serum (FBS) (Gibco), 2 mM L-Glutamine (Invitrogen), 50 μg/ml penicillin (Invitrogen) and 50 μg/mL streptomycin (Invitrogen). HSVECs were obtained by enzymatic collagenase digestion of human saphenous veins (Ethics 15/ES/0094) and maintained in EC growth medium (EGM-2 BulletKit™) (Lonza) supplemented with foetal bovine serum (10%, Life Technologies) and Penicillin-Streptomycin (100U/ml) (Gibco). All cells were used between passages 3-5.

### High-content microscopy high-throughput proliferation screening of miRNAs on human PASMCs

A library of 2042 human miRNAs (Dharmacon, ThermoFisher), was used to transfect human PASMCs at a concentration of 50 nM using RNAiMAX lipofectamine. 1200 cells were plated per well in 384-well plates. 48 h post transfection, cells were pulsed with EdU for 24 h, and fixed 72 h after transfection. Cells were stained with the Alexa fluor 488 EdU Click-it kit (Thermofisher Scientific), and counterstained with Hoechst 33342. High content fluorescent images were acquired using the Image Xpress micro microscope (Molecular Devices) and analysed using MetaXpress version 5.3.0.5 to assess total cell number and percentage of EdU positive cells. MiRNAs exerting a negative effect on cell number, i.e. likely toxic, were excluded from analysis by calculating a z-score of the total cell number count per well and excluding those with a reduction in total cell count of ≥ 1.5 standard deviations compared to the median cell count value.

Four miRNA Controls (miR-CTRL) as well as siRNA targeting Ubiquitin C gene (siUBC) from Dharmacon were included in the screen.

Dharmacon miRNA mimic negative control #1,#2,#3, and #4.

siUBC: siGENOME smartpool Dharmacon (seq#1 GUGAAGACCCUGACUGGUA, seq#2 AAGCAAAGAUCCAGGACAA, seq#3 GAAGAUGGACGCACCCUGU, seq#4 GUAAGACCAUCACUCUCGA).

### Screening validation experiments

The fluorescence-microscopy based analysis of proliferation described above for the screen was used to validate the effect of the 7 selected miRNA mimics (Dharmacon, ThermoFisher) on HPASMCs (n=4), HCASMCs (n=3) and HUVSMCs (n=3). Similar to the screen, this was done in a 384-well plate format using RNAimax lipofectamine and 50 nM miRNA. To monitor proliferation rates, EdU was added to cells after 48 h and fixed after 72 h. Cells were stained, imaged and analysed as mentioned above. miR-CTRL-2 (miR-CTRL-2: cel-miR-239b miRIDIAN microRNA Mimic Negative Control Dharmacon #2 UUGUACUACACAAAAGUACUG) was used as a negative control.

### microRNA (miRNA) mimic mediated transfection of HSVSMC and HSVEC

Transient transfection of the 7 miRNA mimics (Dharmacon, ThermoFisher) or the miRNA control (miR-CTRL-1: cel-miR67 miRIDIAN microRNA Mimic Negative Control #1 UCACAACCUCCUAGAAAGAGUAGA) was performed with Lipofectamine RNAiMAX (Life Technologies) following the manufacturer’s guidelines, for 6 hours in Opti-MEM (Life Technologies).

### RNA extraction, reverse transcription and TaqMan qPCR Analysis

Total RNA isolation was performed using QIAzol Lysis Reagent and the miRNEasy Mini Kit including the RNAse-free DNase Set (Qiagen, Hilden, Germany), according to the manufacturer’s instructions. All RNA samples were stored at −80°C until required. miRNA reverse transcription (RT) to cDNA utilized the Applied Biosystems miRNA Reverse Transcription Kit. cDNA was synthesised from total RNA (2ng/µl RNA per reaction) using the MultiScribe™ Reverse Transcriptase kit (Life Technologies, Paisley, UK) and miRNA-specific RT probes. Thermal cycling conditions for synthesis involved 30-minute incubations at 16°C and 42°C, followed by a 5-minute denaturation at 85°C. The reaction concluded with an indefinite hold at 4°C. After synthesis, all samples were stored at −20°C until required. Target dependant, quantitative real-time polymerase chain reaction (qRT-PCR) was later performed using TaqMan® (Thermo Fisher, Paisley, UK) gene expression assays. TaqMan® qRT-PCR was performed using available TaqMan® Gene Expression probes (table 1 below) following the manufacturer’s protocol (Thermo Fisher, Paisley, UK). RNU48 was used for the normalisation of RT-qPCR. Undetermined Ct values were replaced by 40, the number of qPCR cycles for a typical qPCR run. Quantification of gene expression was analysed as a relative change of gene expression using the 2(-Delta Delta Ct) method as previously described ^36^.

**Table 1:** miRNA TaqMan probes used for qRT-PCR analysis.

| Species | miRNA ID | TaqMan Assay ID |
| --- | --- | --- |
| Human | hsa-miR-323-3p | 002227 |
| Human | hsa-miR-1827 | 002814 |
| Human | hsa-miR-5681b | 476577_mat |
| Human | hsa-miR-4774-3p | 462836_mat |
| Human | hsa-miR-491-3p | 002360 |
| Human | hsa-miR-449b | 001608 |
| Human | hsa-miR-892b | 002214 |
| Human | RNU48 | 001006 |

### Assessment of proliferation using EdU incorporation followed by flow cytometry

Proliferation was assessed using a DNA EdU incorporation assay on a 6-well plate format.

HSVSMC were quiesced in 0.2% FBS media for 48 h prior to transfection. 6h after miRNA mimic or miRNA control transfection, cells were treated with 10 ng/mL IL1α and 20 ng/mL PDGF-BB for 48 h in the presence of EdU. HSVEC were quiesced in 0.2% FBS media for 12 h prior to transfection. 6h after miRNA mimic or miRNA control transfection, cells were cultured in 10% FBS for 48 h in the presence of EdU. Single-cell suspensions of HSVSMCs and HSVECs were fixed for a minimum of 1 h in 70% Ethanol. EdU incorporation was quantified using Click-it EdU Proliferation assay with an Alexa Fluor 488 antibody according to the manufacturer’s protocol (Life Technologies). Cells were resuspended in PBS and analysed by flow cytometry on the BD LSR5 Fortessa Analytic Flow Cytometer using a minimum of 10’000 events. Gating was performed using FlowJo software. One replicate of HSVSMCs transfected with miR-4774-3p was excluded from our analysis as the minimum number of events in flow cytometry was not obtained.

### Wound healing assay

HSVSMCs were plated in 6 well plates and quiesced in 0.2% FBS media for 48 h prior to wound induction and stimulation. Cells were next transfected with 50 nM of miRNA mimic or miRNA control for 6 hours, before scratch with a sterile 1ml pipette tip. Subsequently, scratched HSVSMCs were stimulated with 10 ng/mL IL1α and 20 ng/mL PDGF for 48 h. Post-stimulation, brightfield microscopy images were taken for quantification of wound area closure using ImageJ.

### Caspase-3 assay for apoptosis in HSVSMCs

Caspase-3 activity in HSVSMC lysates was quantified colorimetrically using the Caspase-3 Assay Kit from Abcam (ab39401), according to the manufacturers protocol. Measurement was done on cell lysates with the same total protein concentration. Absorbance at 405nm was measured using a Molecular Devices microplate reader. Cells treated with MG-115 at 1µM concentration were used as a positive control.

### Senescence-associated beta-galactosidase assay in HSVSMCs

Senescence-associated beta-galactosidase in HSVSMC lysates was quantified using the Senescence β-Galactosidase Activity Assay Kit (Fluorescence, Plate-Based) (Cell signaling Technology). Measurement was done on cell lysates with the same total protein concentration. Absorbance at 405nm was measured using a Molecular Devices microplate reader. Cells treated with bleomycin (1µg/mL) were used as a positive control for senescence induction.

### Statistical analysis of *in vitro* results

Biological replicates for HPASMCs, HCASMCs, HUVSMCs correspond to experiments performed on cells derived from different vials. Biological replicates for primary cells HSVSMC and HSVEC correspond to experiments performed using distinct patient derived cell lines. Data are expressed as bar charts of mean ± standard error of the mean (SEM) with individual datapoints superimposed to show full data distribution. Statistical tests used to assess statistical significance is indicated in each figure legend. GraphPad Prism version 10 was used for statistical analysis. For RT-qPCR, statistical analysis was performed on delta Ct, under the assumption of lognormality as previously described ^37^, using a repeated measures ANOVA with Dunnet’s corrections. For EdU incorporation data determined by flow cytometry in HSVSMC, we know that the data follow a normal distribution based on a Shapiro-Wilk test on internal data (data not shown). As the number of replicates is different across conditions, the statistical analysis was done using a mixed-effects model with Dunnet’s corrections. For the other experimental data, as the sample size is <5, normal distribution cannot be assessed accurately. Therefore, data were subjected to Iman-Conover non-parametric ranking followed by Repeated Measures one way ANOVA.

### Analysis of bulk RNA-sequencing following miRNA overexpression

RNAseq was performed on RNA extracted and DNAse treated samples using the miRNeasy mini kit (Qiagen) obtained from three replicates of miRNA overexpression, including controls, in HSVSMCs and HSVECs. Library preparation was prepared after polyA mRNA selection using NEBNext Ultra II Directional RNA Library Prep Kit for Illumina following manufacturer’s instructions (NEB). Sequencing was performed on Illumina NovSeq600 using a 2×150 paired end configuration. Gene quantification (read count and FPKM) was obtained using RSEM (options: -bowtie2 -pairedend), based on human GRCh38 genome annotation and GENCODE transcriptome annotation (Release 25). Raw and processed data of the RNAseq performed in HSVSMCs and HSVECs are available at GEO database under accession Serie GSE253004 with subseries GSE253003 for HSVSMCs data and GSE253002 for the HSVECs data. Downstream analysis was done in RStudio version 2022.07.2 with R version 4.2.2. Principal component analysis and differential expression was performed utilizing DEseq2 version 1.38.3. Removal of batch/patient effect was performed using limma version 3.54.2 removeBatchEffect function. Significantly differentially expressed genes were identified using the following thresholds: absolute Fold Change >=2, adjusted p value of 0.01 and a minimum expression of 2 FPKM in at least 2 of the miR-CTRL/miR-mimic samples. Gene Ontology enrichment analysis was performed using using TopGo version 2.50.0 with Org.Hs.eg.db_3.16.0. Visualisation of genes involved in the KEGG cell cycle pathway hsa04110 was obtained using Pathview version 1.38.0.

### multimiR target prediction and filtering

miRNA target prediction was performed using the multimiR R package. Differentially expressed genes following overexpression of each miRNA were considered targets if they were found among the top 50% of targets (based on prediction score) in two miRNA prediction tools.

### Network analysis and visualisation

Genes commonly regulated by all miRNAs and candidate targets of each miRNA were considered. Genes involved in cell cycle regulation were extracted based on the GO Term annotation GO:0007049. Network analysis was done on these 176 genes. Gene interactions were obtained using STRING version 12.0 (https://string-db.org/) with default parameters. Visualisation of the network was performed using Cytoscape version 3.9.1. Genes commonly regulated by all miRNAs were placed at the centre of the network while targets were placed at the periphery.

## DATA AVAILABILITY STATEMENT

Raw and processed data from the RNAseq has been deposited to GEO database (Serie GSE253004 with subseries GSE253003 for HSVSMCs data and GSE253002 for the HSVECs data). Other data can be obtained upon request.

## ACKNOWLEDGEMENTS

We would like to thank Jean Iyinikkel for conducting experiments in human saphenous vein cell lines and her contribution to the analysis of these experiments and the RNAseq. Flow cytometry data was generated with support from the QMRI Flow Cytometry and cell sorting facility, University of Edinburgh. Bioinformatics analysis has made use of the resources provided by the Edinburgh Compute and Data Facility (http://www.ecdf.ed.ac.uk/).

J.R, A.B and A.H.B are supported by the BHF Chair of Translational Cardiovascular Sciences (CH/11/2/28733). SZ is supported by Grant AIRC IG2020 Id24529. A.H.B is also supported by the BHF program grant (RG/20/5/34796) as well as the ERC Advanced Grant Vascmir (338991).

## AUTHOR CONTRIBUTIONS

AHB, MG, SZ proposed the hypothesis and supervised the study. LB and NARR performed the screen and NARR performed the validation experiments. The experiments in human saphenous vein cell lines were conducted with help from MDB and VM, and the analysis of the experiments were performed by JR, EK, with the help of DK. RNAseq analysis were conducted by JR and EK with the help of DK. JR wrote the manuscript with help of EK. MDB, FV, KM, MB, AB contributed to the discussion. All authors reviewed the manuscripts.

## DECLARATION OF INTERESTS STATEMENT

The authors declare the following financial interests/personal relationships which may be considered potential competing interests: AHB, MG, SZ. are named inventors on a patent application related to this work (PCT/GB2023/052170).

