## Supplemental Figures for "Functional screening identifies novel miRNAs inhibiting Vascular Smooth Muscle Cell proliferation"

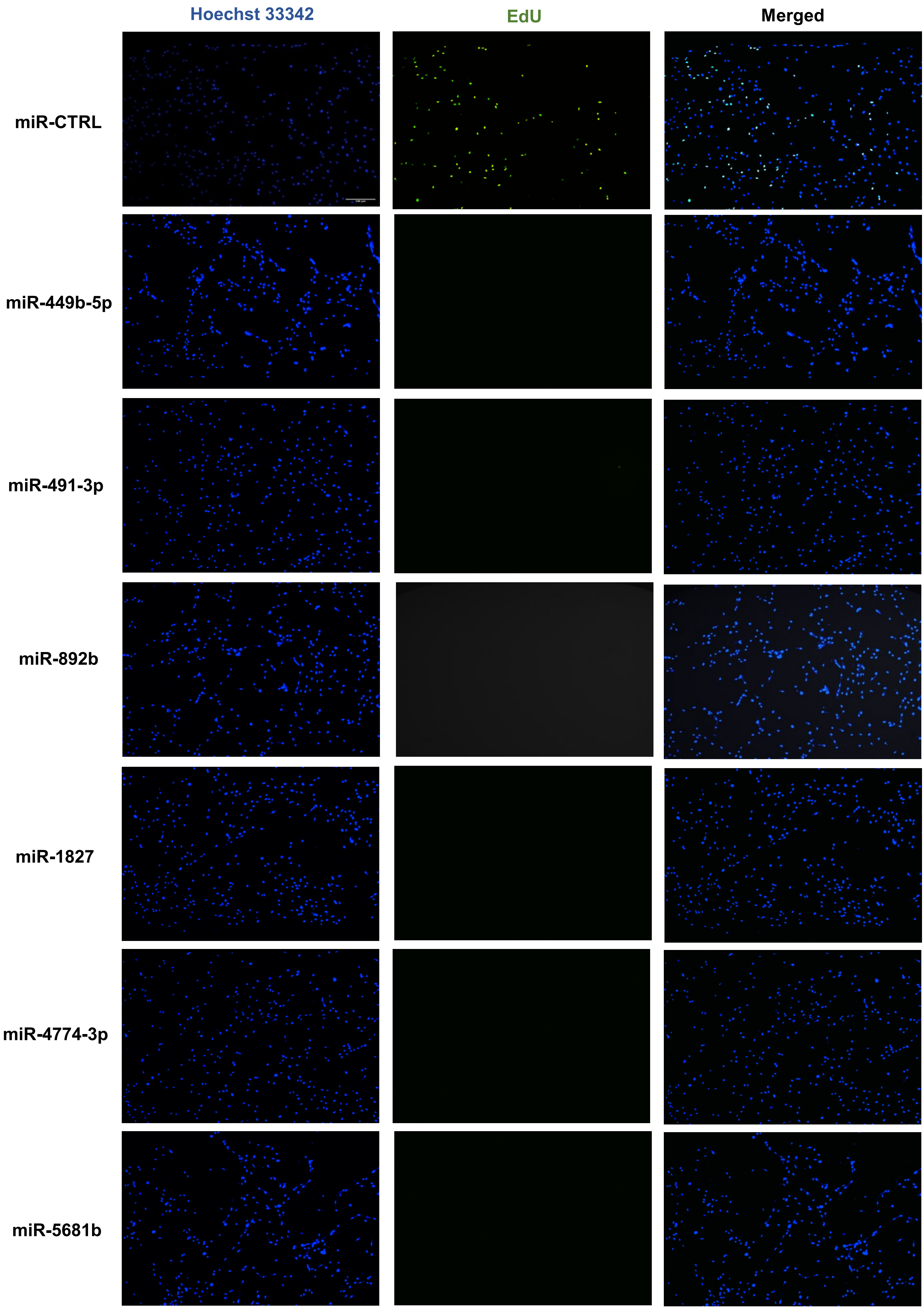


**Supplementary Figure 1.** Representative images of human HPASMCs stained with Hoechst 33342 (blue) and EdU (green) following treatment with miR-CTRL or mimics for miR-449b-5p, miR-491-3p, miR-892b, miR-1827, miR-4774-3p and miR-5681b. Scale bar is 100μm.

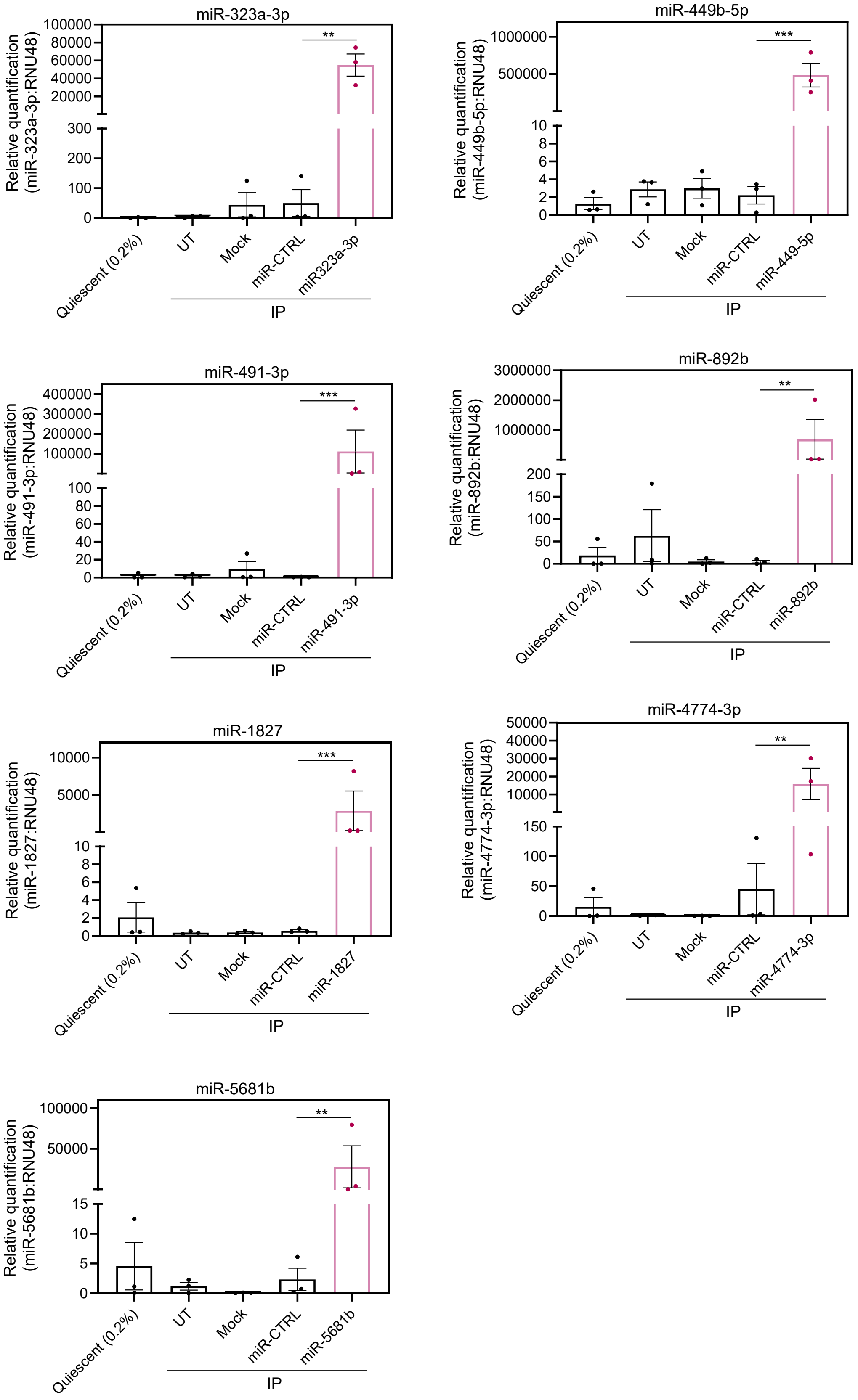


**Supplementary Figure 2**: Confirmation of miRNA overexpression using mimic in HSVSMC by RT-qPCR (n=3). UT: Untransfected, IP: IL1a+PDGF-BB.
Statistical analysis was performed using a repeated measures ANOVA. P-values for the comparison between miRNA mimic treatment and miR-CTRL treatment obtained after Dunnet's test for multiple corrections are included on the graph: **P* < 0.05, ***P* < 0.01, ****P* < 0.001.

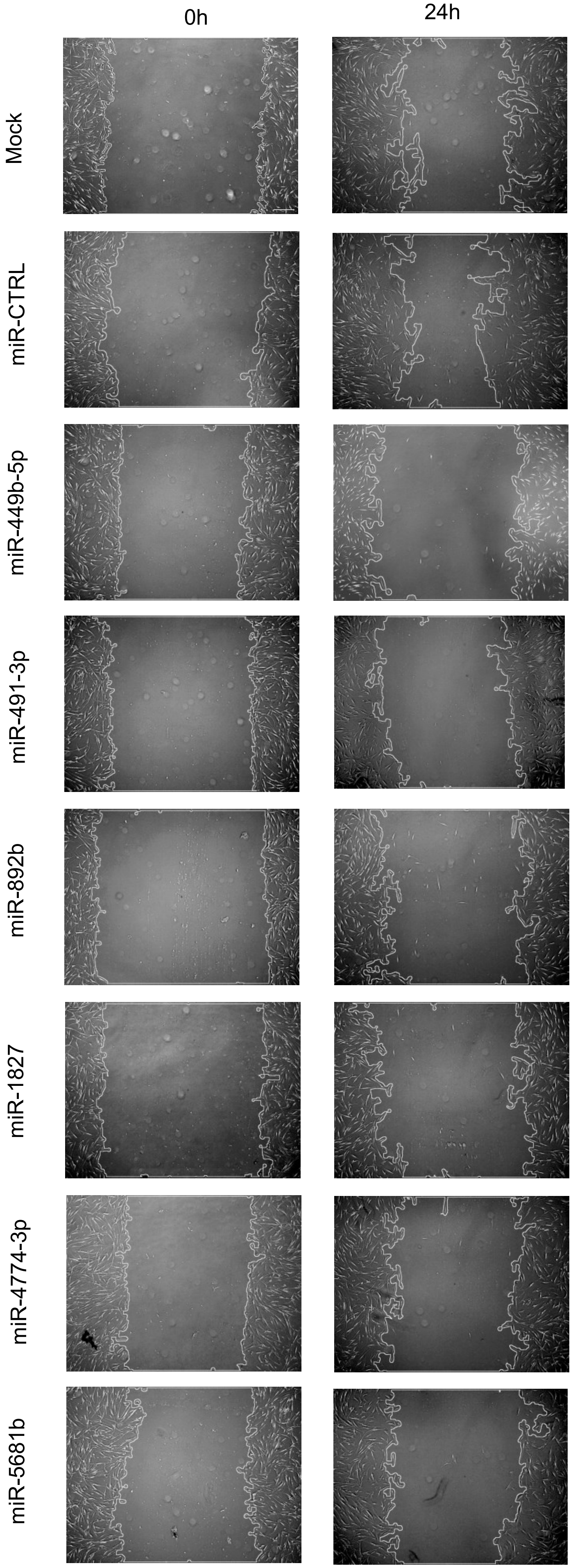


**Supplementary Figure 3**: Wound healing assay of IL-1α/PDGF-BB-stimulated HSVSMCs transfected with the 7 miRNA mimics or miR-CTRL. Representative images from scratch assay at 0 and 24hours for miR-CTRL, miR-449b-5p, miR-491-3p, miR-892b, miR-1827, miR-4774-3p and miR-5681b. Scale bar is 500μm.

A

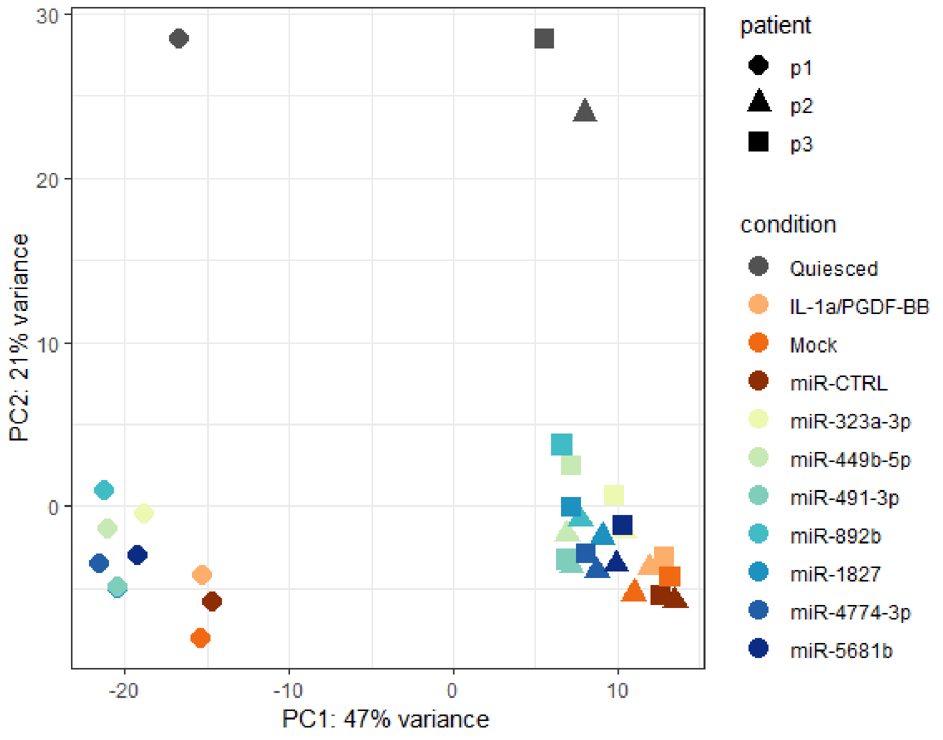


B

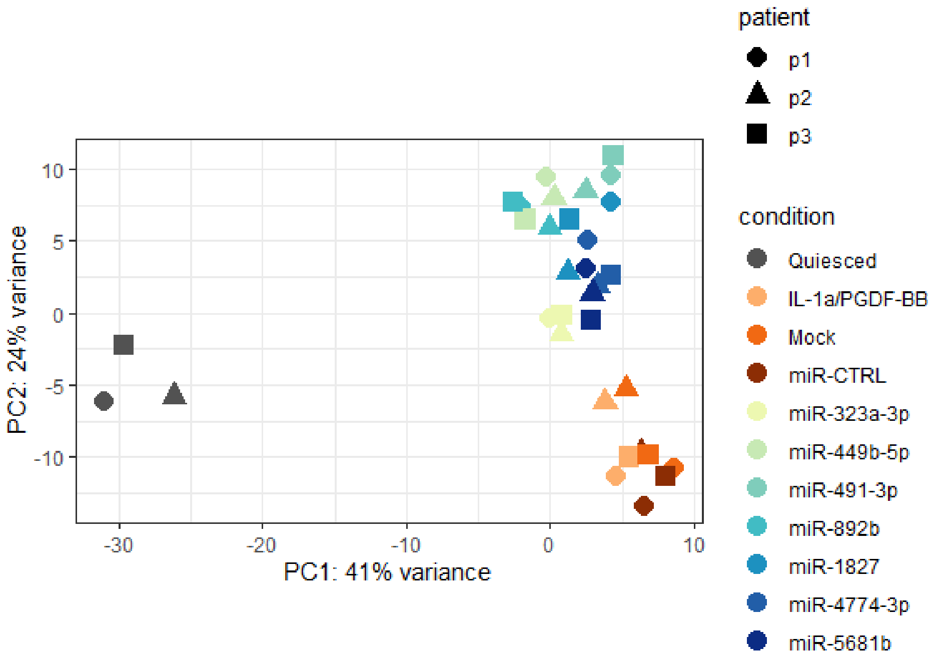


**Supplementary Figure 4**: Principal component analysis plots of the HSVSMC RNAseq without **(A)** and with **(B)** removal of batch effect (removal of patient effect).

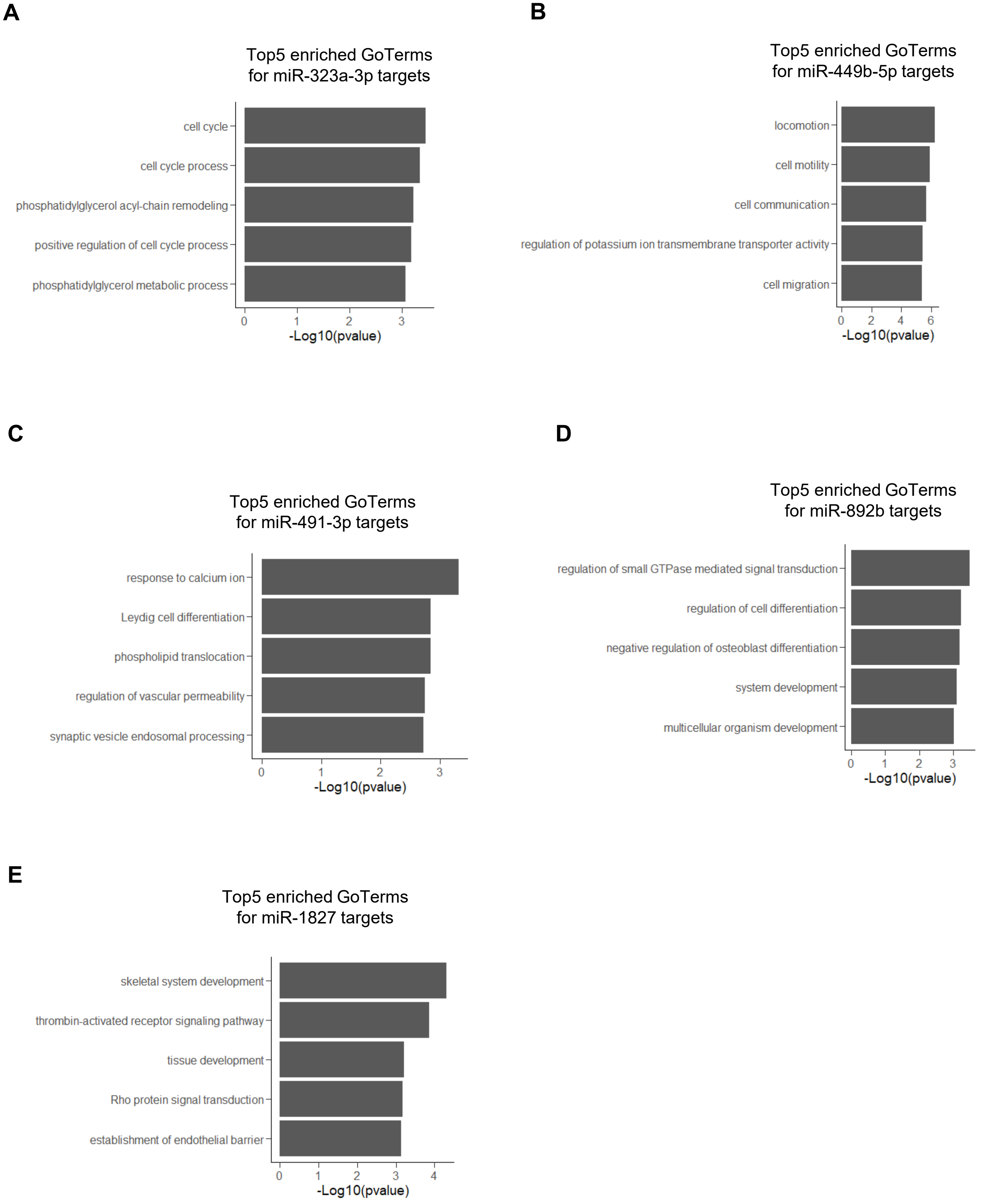


**Supplementary Figure 5**: Top5 enriched Go Terms for the targets of **(A)** miR-323a-3p, **(B)** miR-449b-5p, **(C)** miR-491-3p, **(D)** miR-892b, **(E)** miR-1827.

**A**

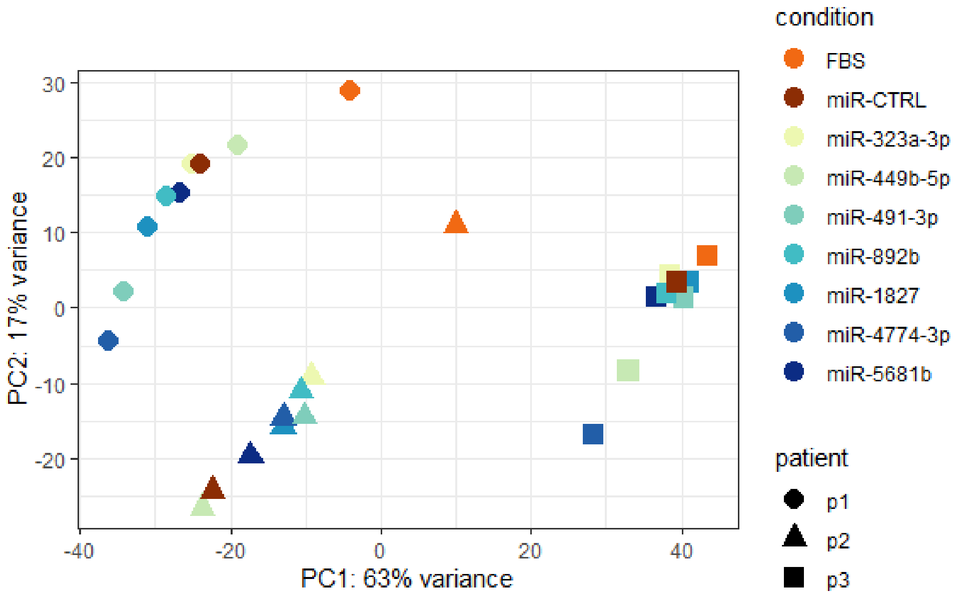


**B**

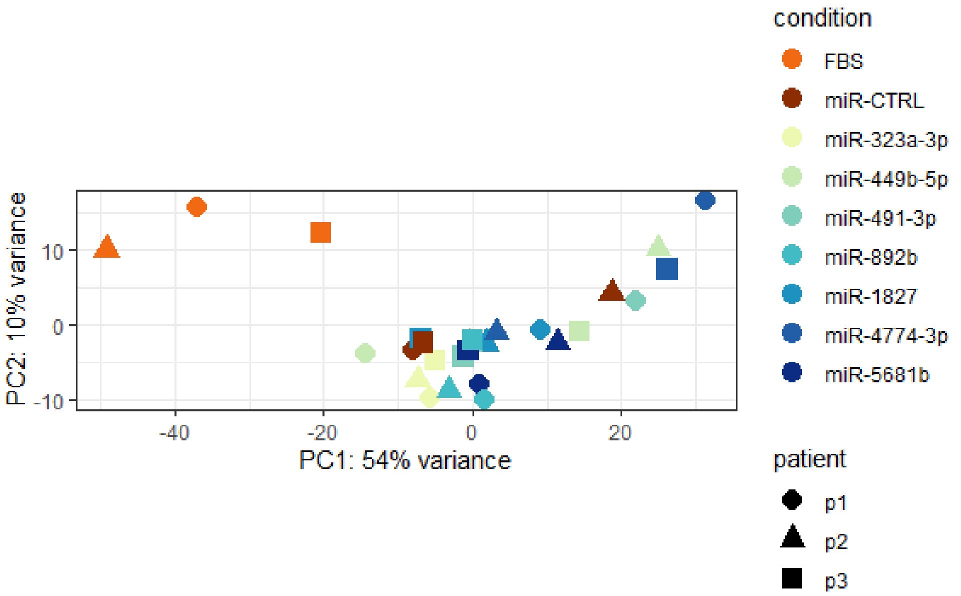


**Supplementary Figure 6:** Principal component analysis plot of the HSVEC RNAseq without **(A)** and with **(B)** removal batch effect (removal of patient effect).

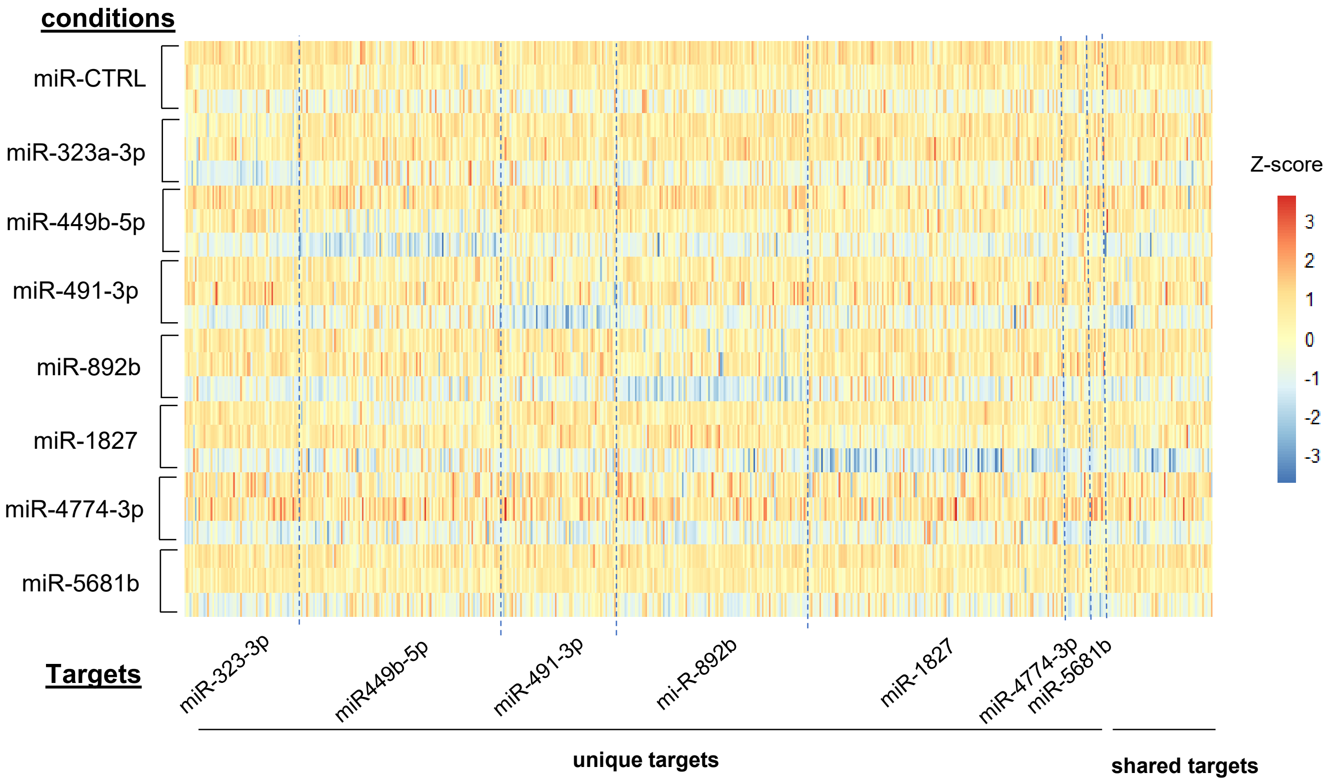


**Supplementary Figure 7:** Heatmap showing the expression profile in HSVEC of all miRNA candidate targets identified in the HSVSMC RNAseq with a separation between unique and shared targets.
